# Age differences in spatial memory are mitigated during naturalistic navigation

**DOI:** 10.1101/2023.01.23.525279

**Authors:** Paul F. Hill, Skyelynn Bermudez, Andrew S. McAvan, Joshua D. Garren, Matthew D. Grilli, Carol A. Barnes, Arne D. Ekstrom

**Author notes:** Correspondence concerning this article should be addressed to Paul Hill or Arne Ekstrom.

## Abstract

Spatial navigation deficits in older adults are well documented. These findings are often based on experimental paradigms that require using a joystick or keyboard to navigate a virtual desktop environment. In the present study, we investigated whether age differences in spatial memory are attenuated when tested in a more naturalistic and ambulatory virtual environment. In Experiment 1, cognitively normal young and older adults navigated a virtual variant of the Morris Water Maze task in each of two virtual reality (VR) conditions: a desktop VR condition which required using a mouse and keyboard to navigate and an immersive and ambulatory VR condition which permitted unrestricted locomotion. In Experiment 2, we examined whether age- and VR-related differences in spatial performance were affected by the inclusion of additional spatial cues in an independent sample of young and older adults. In both experiments, older adults navigated to target locations less precisely than did younger individuals in the desktop condition, replicating numerous prior studies. These age differences were significantly attenuated, however, when tested in the fully immersive and ambulatory environment. These findings underscore the importance of developing naturalistic and ecologically valid measures of spatial memory and navigation, especially when performing cross-sectional studies of cognitive aging.

---

Declines in spatial navigation as part of healthy aging are well documented, suggesting a strong relationship between advanced age and an impaired ability to remember spatial locations. One commonly used paradigm involves virtual adaptations of the Morris water maze (vMWM) task. In these paradigms, older adults typically show significantly reduced accuracy in remembering the location of a hidden target using distal cues compared to younger adults. One reason for the widespread use of this task to assess age-related differences in navigation relates to the ease of administration of the vMWM using desktop computers. The findings from these studies have played a significant role in shaping current models of aging and spatial navigation (Coughlan et al., 2018; Lester et al., 2017; Moffat, 2009).

Studies involving the vMWM task typically require using a keyboard or joystick to navigate a two-dimensional virtual environment on a computer monitor. Results obtained from these types of ‘desktop-based’ virtual reality (VR) studies may be confounded by age differences in prior experience with computer gaming equipment and fine motor control related to joystick and keyboard use. Specifically, older adults may be at a comparative disadvantage due to less exposure to computers and desktop VR compared to younger cohorts (Charness & Boot, 2022). Even studies that have attempted to equate for lower exposure to computer technology in older adults by adding training have still found that comfort ratings with desktop VR are lower in older adults compared to younger adults (Head & Isom, 2010). These findings raise the possibility that age differences in spatial navigation may be exaggerated in less familiar virtual desktop testing situations. Given its frequent use in the field of aging and navigation (Daugherty & Raz, 2017; Daugherty et al., 2015; Iaria, Palermo, Committeri, & Barton, 2009; McAvan et al., 2021; Moffat, Kennedy, Rodrigue, & Raz, 2007; Moffat & Resnick, 2002; Zhong et al., 2017), we focused here on the vMWM task. We note that the vMWM is similar to other tasks frequently tested with older adults as it involves desktop virtual reality and remembering locations using distal landmarks.

Experimental paradigms that use desktop VR rely primarily on visual input and lack the types of naturalistic movements and self-generated idiothetic cues (e.g., motor, proprioceptive, vestibular) that can help form complex spatial representations of the environment (Hejtmanek et al., 2020; Richardson et al., 1999; Ruddle & Lessels, 2006). The absence of these types of body-based cues may disproportionately affect spatial performance in older adults. In an early study by Allen and colleagues (2004), older adults were observed to make more errors on a path integration task compared to younger individuals during trials in which only visual cues were available to navigate (Allen et al., 2004). Young and older adults performed similarly on trials in which proprioceptive and vestibular sensory feedback was available, suggesting that spatial performance might be affected by the presence of multimodal sensory information (see also Adamo et al., 2012). Yu and colleagues (2021) found that, compared to younger individuals, middle-aged adults were significantly impaired in their wayfinding abilities and took fewer shortcuts when navigating a desktop-VR maze (Yu et al., 2021). The two age groups did not differ, however, on an immersive path integration task which, unlike the desktop-VR maze, provided both visual and proprioceptive feedback. Notably, these experiments focused specifically on path integration, which is particularly reliant on precise self-motion feedback.

The overarching aim of the current study was to test the hypothesis that age-related differences in spatial memory may be mitigated in more ecologically enriched test situations. Alternatively, performance on the vMWM may be impaired in older relative to younger adults regardless of the availability of self-motion cues and the more immersive visual experience available with head-mounted displays, suggesting a general age-related impairment on the task. To address these possibilities, we performed two experiments comparing spatial memory performance in cognitively healthy young and older adult humans as they navigated a virtual MWM adapted from McAvan et al. (2021) in each of two conditions: 1) a desktop VR condition that required using a keyboard and mouse to navigate, and 2) an immersive and ambulatory VR condition in which participants wore a wireless head-mounted display that permitted unrestricted locomotion and self-generated idiothetic feedback during navigation. Critically, this study differed from McAvan et al., (2021) in that all participants were tested in both the desktop and immersive virtual environments to permit within-subject comparisons between the respective VR testing conditions.

## Materials & Methods

### Participants

Young and older adult participants were recruited from the University of Arizona and surrounding communities. All participants gave informed consent in accordance with the University of Arizona Institutional Review Boards and were compensated at the rate of $18 per hour. All participants had normal or corrected-to normal color vision, normal or corrected-to-normal hearing, and reported no history of cardiovascular problems, neurological or psychiatric conditions, or history of motion sickness.

### Neuropsychological Test Battery

All older adult participants completed a neuropsychological test battery on a day prior to the experimental VR session. The test battery included multiple tests and scores in each of four broad cognitive domains: memory [California Verbal Learning Test-II (CVLT) Long Delay Free Recall (Delis, DC et al., 2000), Rey-Osterrieth Complex Figure Test Long Delay Free Recall (RCFT-LDFR) (Rey, 1941)], executive function [Trail Making Tests A and B total time] (Reitan, RM & Wolfson, D, 1985), language [F-A-S fluency (Spreen, O & Benton, AL, 1977), category fluency (Benton, 1968), Boston Naming Test (BNT); Goodglass et al., 2001], visuo-spatial abilities [Wechsler Adult Intelligence Scale 4^th^ edition (WAIS-IV) Block Design test (Wechsler, D, 2009), RCFT copy score (Rey, 1941)], and verbal intelligence [WAIS-IV Vocabulary and Similarities] (Wechsler, D, 2009). Participants were excluded from entry into the study if they scored < 1.5 SDs below age-appropriate norms on any memory test or < 1.5 SDs below age norms on any two other tests. These criteria were employed to minimize the likelihood of including older individuals with mild cognitive impairment, individuals who are considered at elevated risk for Alzheimer’s disease and related dementias. Results from the neuropsychological test battery are presented in Supplementary Table 1.

### Virtual Environments

The virtual environment and experimental tasks were built in Unity 3D (Unity Technologies ApS, San Francisco, CA) using the Landmarks virtual reality navigation package (Starrett et al., 2021). The navigable virtual environment was approximately 5×5 m in size, with the full space spanning 750×750 m. Four distally rendered mountains were visible from within the 5×5 m space. Each virtual environment was rendered with a unique floor (snow-covered or desert), and three unique 3D rendered objects (book, puzzle cube, teapot in the snow-covered environment; alarm clock, mug, rotary phone in the desert environment). These objects were presented on pedestals at approximately chest height and served as the hidden navigation goals (see Procedures). The respective virtual environments and order of administration were fully counterbalanced across the immersive and desktop VR conditions.

#### Immersive VR Condition

To simulate the immersive experience of being in a mountainous environment, we used the HTC Vive Pro headmounted display (HMD) in conjunction with the HTC wireless Adapter (HTC, New Taipei City, Taiwan) to allow for untethered, free ambulation. The Vive Pro displayed stimuli at a resolution of 1140-1600 pixels per eye, 90 Hz refresh rate, and a 110_J field of view, while the Wireless Adapter delivered data over a 60 GHz radio frequency for up to 7m. Participants were allowed to interact with the virtual environment and record their responses using a handheld HTC Vive controller (HTC, New Taipei City, Taiwan). The immersive VR task was run on a custom-build computer with a NVIDIA GeForce Titan Xp graphics card (NVIDIA Corp., Santa Clara, CA, United States.

#### Desktop VR Condition

All participants navigated an analogous version of the MWM task optimized for a laptop computer. The desktop VR condition was completed in a quiet behavioral testing room with participants seated approximately 2’ from the screen. The task was run on a 15” Lenovo Legion Y540 gaming laptop computer with a GTX1660 Ti graphics card. Forward, left, backwards, and right translations were made by pressing the ‘W’, ‘A’, ‘S’, and ‘D’ keys, respectively. Participants could simultaneously use the mouse to rotate their view around the xyz axis.

### Navigation Procedures

Prior to beginning the immersive phase of the experiment, participants were instructed to close their eyes as they were led into the navigation room. This prevented participants from seeing the size and shape of the physical environment and heightened the level of immersiveness. Upon entering the navigation room, participants were fitted with the wireless HMD, a handheld controller (held in their dominant hand), and a clip-on battery pack that powered the wireless HMD. Participants were then immersed in a practice virtual environment similar to the main task described here. During the practice phase, participants were allowed to freely navigate a small circular room for five trials lasting 30 seconds before being prompted to find and remember the location of a single target object for a subsequent five trials. The practice session lasted approximately 10 min. Participants completed an identical practice session prior to beginning the desktop condition to ensure proficiency with using the keyboard and mouse to navigate.

After the practice session in each VR condition, participants were tasked with completing five blocks of a navigation task. Participants were offered an opportunity to take brief breaks in-between each block. The study design was based on a prior study conducted in healthy young and older adults performed in an immersive VR environment (McAvan et al., 2021). A similar version of this task was also used with amnestic patients in a desktop VR environment (Kolarik et al., 2018). In each task block, participants received verbal and visual instructions and were provided with reminders when requested. White noise was played through headphones throughout the experiment to prevent sound cues from providing location or orientation information.

After completion of each trial, participants were briefly disoriented by guiding them around the environment without vision. This prevented participants from tracking their bearing and movements through the environment from trial to trial and thus ensuring participants used their memory to find the target locations. During the immersive VR condition, participants were physically guided by a research assistant along a random path while the HMD presented a blank screen. During the desktop VR condition, participants viewed a blank screen during the disorientation period, after which they were placed in the next start location. The order and sequence of the start locations were held constant for each virtual environment across all participants.

#### Spatial Learning Block

Participants were familiarized with the locations of the six target objects (three in each VR environment) by performing 16 blocked spatial learning trials. Before each trial, participants were disoriented for 30 s and then placed at one of eight starting positions. Each spatial learning block was organized into four sub-blocks, each corresponding to a unique start position (e.g., 4 consecutive trials from position 1, followed by 4 consecutive trials from position 5, etc.). Each trial began with visual instructions indicating the navigation goal (e.g., ‘Please find the book’). Participants then freely navigated the environment until the target object appeared after 30 s.

After the first trial of each learning block, participants were encouraged to remember the location of the target object and to walk to that location before 30 s elapsed. Participants had the option to make a button response if they were confident they had navigated to the correct target location. This was recorded as a spatial memory response and the location and timestamp of this response were logged in the data output. Upon making a button press, the target object became visible and participants were instructed to touch the object with the handheld controller to move to the next trial. If a button response was not made, the target object would appear after 30 s, at which point the location and timestamp were logged in the data output and participants were permitted to touch the object to move to the next trial.

#### Immediate Spatial Probe Trials

After 16 trials of each spatial learning block, participants performed a single probe trial. Participants were placed in a novel start position and tasked with recalling the location of the target object. Participants were instructed that the object would not appear after 30 s, nor would the object appear after making a button press. Note that this differed from McAvan et al. (2021) in which the object appeared immediately after the button press. The trial commenced with visual instructions indicating the navigation goal (e.g., ‘Please find the book’). Upon reaching the location of the hidden target object, participants made a button press and their location and timestamp was logged in the data output. Participants then had the opportunity to take a brief break before moving onto the next block.

#### Visual Target Block

Following the three spatial learning blocks, participants completed one block of eight visible target trials. The target objects remained visible for the entire duration of these trials. These trials served as a control for motivational and potential sensorimotor deficits in performing the task (Moffat & Resnick, 2002; Morris et al., 1982), although they have also been employed as a comparison for “egocentric” navigation. Before each trial, participants were disoriented for 20 s while the virtual environment was blacked out. Each trial began with visual instructions indicating the navigation goal (e.g., ‘Please find the book’), at which point the participants simply walked to the visible object and touched it with the handheld controller in order to progress to the next trial. Each target object was presented in sequential order (i.e., target 1, target 2, target 3, target 1, target 2, etc.).

#### Delayed Spatial Probe Trials

Following the visible target block, participants performed 12 probe trials designed to test delayed spatial recall. Before each trial, participants were disoriented for 20 s while the virtual environment was blacked out. Each trial began with visual instructions indicating the navigation goal (e.g., ‘Please find the book’). Participants were instructed to find each target object in sequential order (e.g., target 1, target 2, target 3, target 1, target 2, etc.). Six of the 12 delayed probe trials began from a start position that previously accompanied that target object during the spatial learning block (‘repeated’ trials). The remaining six delayed probe trials began from a novel start position that was not previously linked to the target object (‘novel’ trials). As with the immediate probe trials, the target object did not appear after 30 s, nor did it appear after a button press. Again, this differed from McAvan et al. (McAvan et al., 2021) in which the object appeared immediately after the button press. Participants were thus required to navigate to the location of the hidden target object based solely on their memory. Upon reaching this location, participants made a button press to record their location and timestamp.

#### Rotated Mountain Trials

The final block comprised three probe trials that were similar to the immediately preceding delayed probe trials, the sole difference being that one of the distal mountain cues was rotated 20_J clockwise or counterclockwise around the target. The objective was to explore how manipulation of the distal navigation cues affected age differences in navigation strategies. These analyses are beyond the scope of the present report and will not be discussed in any further detail here.

### Statistical Analyses

Position and rotation data was recorded throughout the entire experiment at a sampling rate of approximately 10 Hz. All statistical analyses were conducted with R software (R Core Team, 2017). All *t*-tests were two-tailed and performed using the t.test function in the base R package. Welch’s unequal variance *t*-tests were performed when assumptions of equal variance were not met. ANOVAs were conducted using the *afex* package (Singmann, H. et al., 2016) and the Greenhouse-Geisser procedure (Greenhouse & Geisser, 1959) was used to correct degrees of freedom for non-sphericity when necessary. Post-hoc tests on significant effects from the ANOVAs were conducted using the *emmeans* package (Lenth, R., 2018) and corrected for multiple comparisons using the Holm-Bonferroni procedure where appropriate. Bayes factor values were computed using the *BayesFactor* package (Morey et al., 2022).

Spatial precision was operationalized as the Euclidean distance between the remembered and true location of each target object (i.e., distance error). We computed the mean trial-wise distance error separately for each participant and experimental block (spatial learning trials, immediate probe trials, and delayed probe trials). During the spatial learning block, the *remembered* location corresponded to a participant’s location when a memory response was indicated on the handheld controller/mouse or, in the absence of an overt button response, by recording participant location after 30 seconds had elapsed and the target object became visible. During the immediate and delayed probe trials, the *remembered* location always corresponded to the participant location when a memory response was indicated (as the target objects did not appear after any amount of time). Unless otherwise specified, we report the results of two-way mixed-factorial ANOVAs with factors of age group (young, older) and VR condition (desktop, immersive).

## Experiment 1

In Experiment 1, young and older adult participants were trained to locate hidden targets in each of two virtual MWM environments: a desktop VR condition which required using a mouse and keyboard to navigate and an immersive and ambulatory VR condition which permitted unrestricted locomotion. Memory was assessed immediately after learning (immediate probe) and after a delay (delayed probe). The principal outcome variable was spatial precision, operationalized as the Euclidean distance between the remembered and actual location of each target object.

### Participants

We recruited 21 young and 22 older adults from the University of Arizona and surrounding communities. One older adult voluntarily withdrew from the study due to a scheduling conflict; one young and one older adult withdrew after experiencing mild motion sickness during the virtual reality task. The final participant sample included 20 young adults (18-27 yrs.; *M* = 21.15 yrs; *SD* = 2.30 yrs; 12 females) and 20 older adults (66-80 yrs.; *M* = 73.30 yrs.; *SD* = 3.89 yrs.; 7 females). The final sample size (N = 40) was informed by an *a priori* power analysis conducted G*Power employing between-repeated measures. The sample was selected to ensure power of 0.85 (at *p* < .05) to detect medium age group x VR condition interactions (*f* =.25) in a 2 (age group) x 2 (VR condition) ANOVA.

### Results

Descriptive statistics for the primary outcome measures are summarized in Table 2. During the initial spatial learning trials (Figure 2, top panel), we observed a significant interaction between age group and VR condition (*F*_(1,38)_ = 11.75, *p* = .001, partial-_η_^2^ = .236, BF_10_ = 20.406). This interaction was driven by significantly greater distance error in the desktop compared to immersive condition in older adults (*t*_(38)_ = 5.71, *p* < .001, BF_10_ = 19108.740), but no significant difference between the respective conditions in young adults (*t*_(38)_ = 0.86, *p* = .394, BF_10_ = 0.297). Mean spatial precision was significantly reduced in older adults relative to younger individuals in both the desktop and immersive VR conditions (*t*_(38)_ = 5.81, *p* < .001, BF_10_ = 11970.360; *t*_(38)_ = 4.32, *p* < .001, BF_10_ = 201.724, respectively).

**Figure 1.**
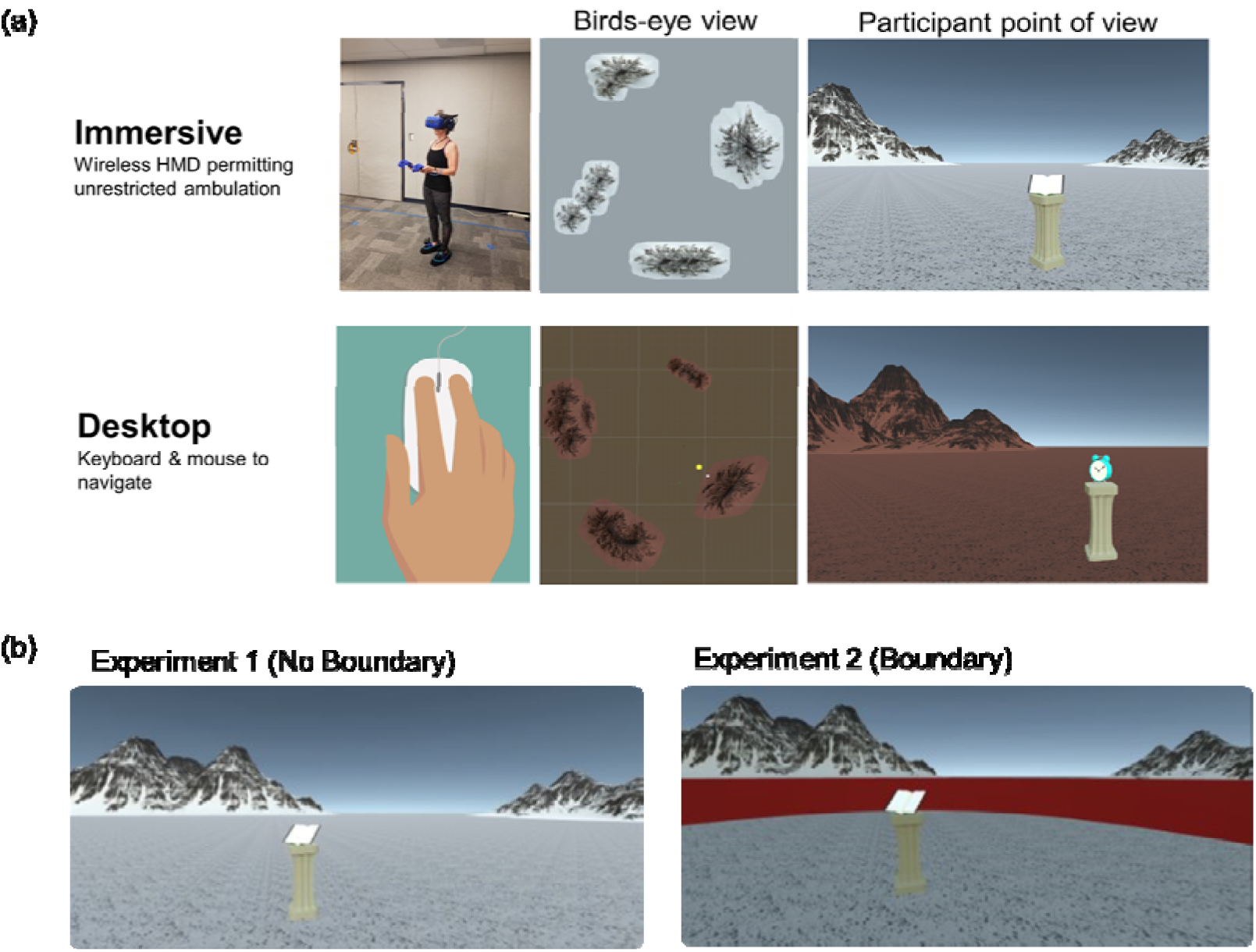
Virtual Morris Water Maze Task Design. (a) Visual depiction of the immersive and desktop VR conditions. The assignment of the respective environments (snow, desert) to each VR condition (desktop, immersive) was fully counterbalanced across participants, as was the testing order of the respective conditions. During the spatial learning blocks, participants were trained to locate the spatial positions of three hidden target objects. Immediately following each spatial learning block participants were cued to recall the location of the hidden object without feedback (immediate probe trials). Following the learning blocks, participants completed eight visible target trials to rule out potential age-related motivational and/or sensorimotor confounds. Participants then completed 12 delayed probe trials without feedback, during which they were cued to locate the hidden target. In a subset of the delayed probe trials, a singular mountain cue was rotated 20 degrees clockwise or counterclockwise unbeknownst to the participant. (b) First person depiction of the available spatial cues in Experiment 1 (distal mountains only) and Experiment 2 (distal mountains and perimeter boundary).

**Figure 2.**
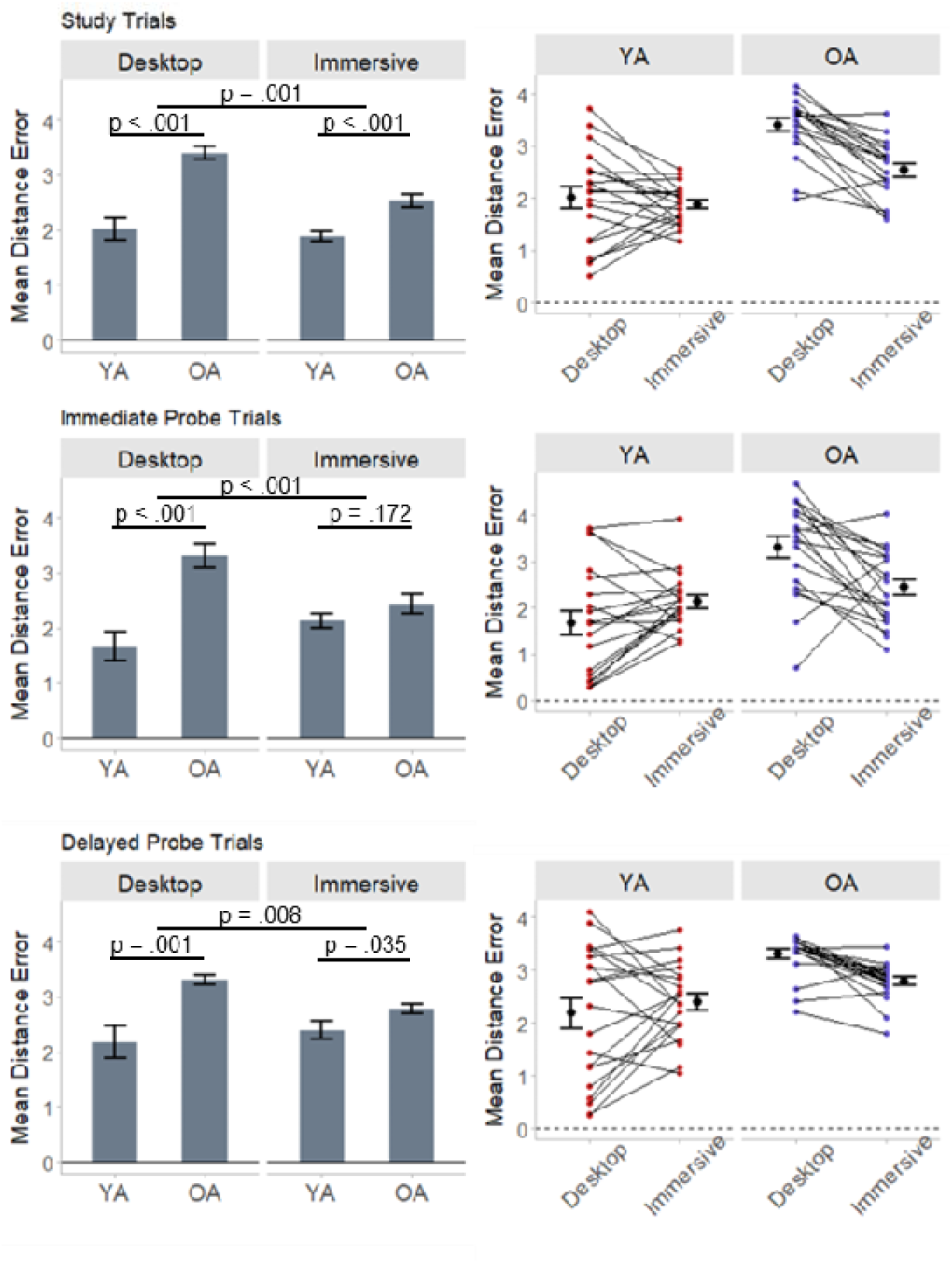
Mean spatial distance error is plotted for the spatial learning trials (top panels), immediate probe trials (middle panels), and delayed probe trials (bottom panels). The spaghetti plots illustrate within-subject changes in distance error between the respective VR conditions. Error bars reflect standard error of the mean.

A similar interaction between age group and VR condition was also evident during the immediate (*F*_(1,38)_ = 15.66, *p* < .001, partial-_η_^2^ = .292, BF_10_ = 117.881; Figure 2, middle panel) and delayed (*F*_(1,38)_ = 7.93, *p* = .008, partial-_η_^2^ = .173, BF_10_ = 6.792; Figure 2, bottom panel) probe trials. In both instances, this interaction was driven by significantly greater distance error in the desktop vs. immersive VR condition in older adults (immediate: *t*_(38)_ = 3.67, *p* = .001, BF_10_ = 12.819; delayed: *t*_(38)_ = 2.82, *p* = .008, BF_10_ = 797.036), along with a null effect of VR condition among young adults (immediate: *t*_(38)_ = -1.92, *p* = .062, BF_10_ = 1.530; delayed: *t*_(38)_ = - 1.16, *p* = .252, BF_10_ = 0.329). Significant age differences in spatial precision were evident in the desktop VR condition (immediate: *t*_(38)_ = 4.71, *p* < .001, BF_10_ = 562.072 ; delayed: *t*_(38)_ = 3.70, *p* = .001, BF_10_ = 42.881). Mean distance error was also significantly greater in older adults compared to young adults in the immersive VR condition during the delayed probe trials (*t*_(38)_ = 2.18, *p* = .035, BF_10_ = 1.933). The two age groups did not significantly differ, however, during the immediate probe trials when navigating in the immersive VR environment (*t*_(38)_ = 1.39, *p* = .172, BF_10_ = 0.663), although the Bayes value suggested only weak or ‘anecdotal’ evidence for null hypothesis.

We next examined the effect of age group and VR condition on the amount of time spent navigating during the immediate and delayed probe trials (Figure 3A), as older adults could be compromising reaction time for accuracy (Salthouse, 1991). For each trial, we computed the log-transformed latency between trial onset and the button response marking spatial memory. We then computed the mean trial-wise navigation latency and submitted these values to a two-way mixed-factorial ANOVA with factors of age group and VR condition. This analysis revealed non-significant main effects of age group (*F*_(1,38)_ = 2.54, *p* = .119, partial-_η_ = .063, BF_10_ = .951) and VR condition (*F*_(1,38)_ = 2.32, *p* = .136, partial-_η_ = .057, BF_10_ = .536) during the immediate probe trials. A significant interaction between age group x VR condition on time spent navigating during the immediate probe trials (*F*_(1,38)_ = 7.52, *p* = .009, partial-_η_ = .165, BF_10_ = 4.881) was driven by younger adults spending more time navigating in the immersive VR condition compared to desktop VR (*t*_(38)_ = -3.02, *p* = .005, BF_10_ = 5.366). Time spent navigating among older adults did not significantly differ between the respective VR environments (*t*_(38)_ = 0.86, *p* = .393, BF_10_ = .334). To examine whether age- and/or VR-related differences in navigation time might moderate commensurate differences in spatial precision, we regressed out the effects of navigation time on distance error and then submitted the resultant residualized distance error values to a two-way ANOVA with factors of age group and VR condition. The interaction between age group and VR condition on distance error remained significant when controlling for navigation time (*F*_(1,38)_ = 22.19, *p* < .001, partial-_η_^2^ = .369, BF_10_ = 1061.301).

**Figure 3.**
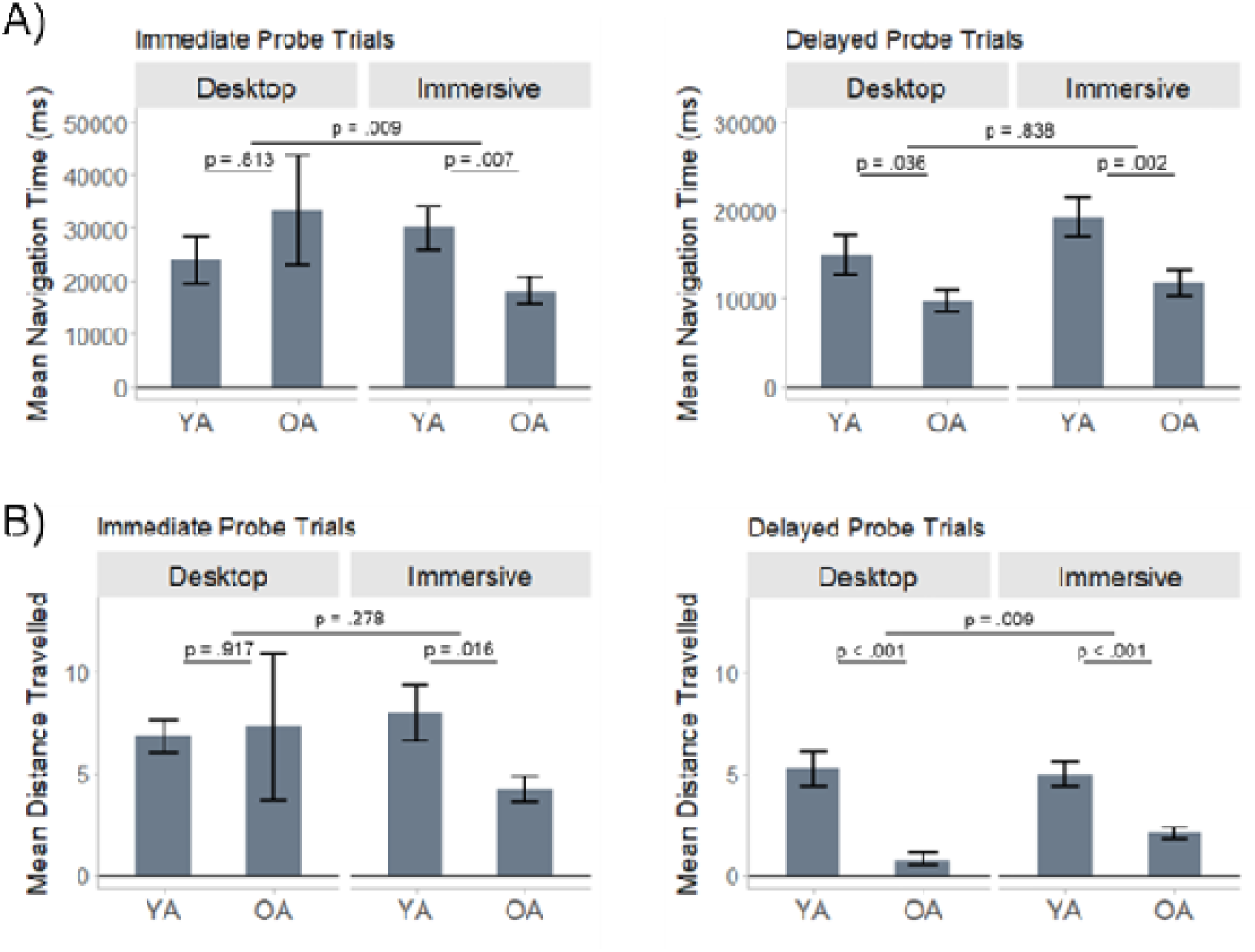
Mean (a) navigation time and (b) total distance travelled is plotted for each age group and VR condition during the immediate probe trials (left panels) and delayed probe trials (right panels). Error bars reflect standard error of the mean.

An identical analysis performed on the delayed probe trials revealed a significant main effect of age group (*F*_(1,38)_ = 9.04, *p* = .005, partial-_η_^2^ = .192; BF_10_ = 9.151) which was driven by *shorter* navigation latencies in older relative to younger adults (*t*_(38)_ = -3.01, *p* = .005; BF_10_ = 52.877), and a significant main effect of VR condition (*F*_(1,38)_ = 11.91, *p* = .001, partial-_η_^2^ = .239; BF_10_ = 24.754) which was driven by shorter navigation latencies in the desktop relative to immersive VR condition (*t*_(38)_ = -3.45, *p* = .001; BF_10_ = 1.949). The interaction between age group and VR condition was not significant (*F*_(1,38)_ = 0.04, *p* = .838, partial-_η_^2^ = .001; BF_10_ = 0.303). A two-way ANOVA of residulized distance errors confirmed that the interaction between age group and VR condition on distance error remained significant when controlling for total time spent navigating (*F*_(1,38)_ = 8.44, *p* = .006, partial-_η_^2^ = .182; BF_10_ = 7.658).

We next computed the total distance travelled during each of the immediate and delayed probe trials and submitted the mean values to a two-way mixed-factorial ANOVA with factors of age and VR condition (Figure 3B). The main effects of age group, VR condition, and the interaction between age and VR condition on total distance traveled during the immediate probe trials were all non-significant (*ps* > .270, BF_10_ < 0.556). A significant interaction between age group and VR condition on total distance travelled was evident during the delayed probe trials (*F*_(1,38)_ = 7.55, *p* = .009, partial-_η_^2^ = .166; BF_10_ = 4.734). This interaction was driven by older adults travelling less distance in the desktop VR condition (*t*_(38)_ = -3.20, *p* = .003, BF_10_ = 161.695) along with a null effect of condition among younger adults (*t*_(38)_ = 0.69, *p* = .496, BF_10_ = 0.267).

To examine whether total distance travelled moderated any of the age and/or VR condition differences in spatial precision, we regressed out the effects of total distance on distance error and then submitted the resultant residualized distance error values to a two-way ANOVA with factors of age group and VR condition. The interaction between age group and VR condition on distance error remained significant when controlling for total distance travelled during the immediate (*F*_(1,38)_ = 18.30, *p* < .001, partial-_η_^2^ = .325; BF_10_ = 302.403) and delayed (*F*_(1,38)_ = 4.34, *p* = .044, partial-_η_^2^ = .102; BF_10_ = 1.640) probe trials, though the Bayes factor indicated only anecdotal evidence in favor of the alternative hypothesis during delayed probe trials.

Following the three spatial learning blocks, participants completed one block of eight visible target trials, during which the targets remained visible for the duration of the trial. These trials were included to rule out potential age-related motivational and/or sensorimotor deficits which may have confounded performance on the task. A 2 (age group) x 2 (VR condition) ANOVA of distance error revealed a significant main effect of condition (*F*_(1,38)_ = 11.29, *p* = .002, partial-_η_ = .229; BF_10_ = 0.920) which was driven by greater distance error in the immersive compared to the desktop VR condition. The main effect of age and the age x condition interaction were not significant (*p*s > .1; BF_10_ < 0.395).

For each participant, we computed the mean log-transformed time it took to reach the visible targets and submitted these values to a 2 (age group) x 2 (VR condition) ANOVA. The analysis revealed a main effect of VR condition (*F*_(1,38)_ = 3.98, *p* = .053, partial-_η_ = .095; BF_10_ = 1.330) which was slightly beyond the *a priori* significance criterion (α < .05). This marginal effect was driven by shorter navigation times in the immersive (*M* = 6972 ms, *SD* = 3546 ms) relative to the desktop (*M* = 9421 ms, *SD* = 9161) VR condition. The main effect of age and the age x condition interaction were not significant (*ps* > .1; BF_10_ < 0.824).

### Discussion

In Experiment 1, we found that age differences in the precision of spatial memories were moderated by the modality of the virtual testing environment. Older adults performed disproportionately worse when attempting to navigate a two-dimensional virtual desktop environment. By contrast, spatial performance in young adults did not reliably differ between the desktop and immersive testing conditions. Control analyses confirmed that the deleterious effects of the desktop VR condition in older adults could not be explained by systematic age- or condition-related differences in total distance travelled, total time spent navigating, or sensorimotor confounds.

## Experiment 2

In the first experiment, participants were required to find the location of a hidden target object relative to four distal landmarks. Our findings suggested that age differences persisted in immersive VR, although these age differences were significantly attenuated compared to desktop VR. One possibility is that older adults were less effective than young adults at using the distal landmark cues to navigate. Recent findings suggest that the ability to orient and navigate with reference to external landmarks declines during healthy aging (Colmant et al., 2023; West et al., 2023), and that older adults may benefit from boundary cues (Bécu et al., 2019; 2023). Therefore, in the second experiment, we included a circular boundary wall located outside the perimeter of the navigable space (diameter = 20 virtual meters). This wall served as a geometric boundary cue that could be used in conjunction with the distal landmarks to orient and estimate distances within the MWM environment. We were specifically interested in whether the inclusion of additional visual cues might affect the age- and VR-related differences in spatial precision that were identified in the first experiment. We predicted that spatial performance would generally be improved by the inclusion of additional boundary cues, but that older adults would still be at a comparative disadvantage compared to young adults when navigating in the virtual desktop environment. We also implemented a mandatory five-minute break between the visible target trials and the delayed probe trials to minimize the possibility that locations were maintained in working memory.

### Participants

Twenty-four young (18-31 yrs; *M* = 21.79 yrs; *SD* = 3.87 yrs; 14 females) and 24 older adults (63-79 yrs; *M* = 68.13 yrs; *SD* = 4.41 yrs; 14 females) were recruited from the University of Arizona and surrounding communities to participate in Experiment 2.

### Results

Descriptive statistics for the primary outcome measures are summarized in Table 2. Mirroring the results of Experiment 1, significant interactions between age group and VR condition on spatial precision were evident during spatial learning trials (*F*_(1,46)_ = 19.88, *p* < .001, partial-_η_^2^ = .302, BF_10_ = 1122.626), immediate probe trials (*F*_(1,46)_ = 9.25, *p* = .004, partial-^η2^ = .167, BF_10_ = 22.988), and delayed probe trials (*F*_(1,46)_ = 13.24, *p* < .001, partial-_η_^2^ = .188, BF_10_ = 89.029). As can be seen in Figure 4, older adults performed reliably worse in the desktop condition compared to the immersive VR condition across all task phases (learning: *t*_(46)_ = 5.09, *p* < .001, BF_10_ = 61.245; immediate: *t*_(46)_ = 2.91, *p* = .006, BF_10_ = 2.427; delayed: *t*_(46)_ = 4.13, *p* < .001, BF_10_ = 55.210). By contrast, spatial precision did not reliably differ between the respective VR conditions in young adults (learning: *t*_(46)_ = -1.22, *p* = .230, BF_10_ = 1.037; immediate: *t*_(46)_ = -1.39, *p* = .170, BF_10_ = 0.910; delayed: *t*_(46)_ = -1.02, *p* = .314, BF_10_ = 0.355). Spatial precision was significantly reduced in older adults compared to young adults in all task phases of the desktop VR condition (learning: *t*_(46)_ = 6.47, *p* < .001, BF_10_ = 176928.700; immediate: *t*_(46)_ = 4.81, *p* < .001, BF_10_ = 1048.590; delayed: *t*_(46)_ = 6.33, *p* < .001, BF_10_ = 112677.100). Age-related effect sizes were smaller in the immersive VR condition but remained significant across all task phases (learning: *t*_(46)_ = 4.12, *p* < .001, BF_10_ = 147.466; immediate: *t*_(46)_ = 3.18, *p* = .003, BF_10_ = 14.086; delayed: *t*_(46)_ = 4.14, *p* < .001, BF_10_ = 156.210).

**Figure 4.**
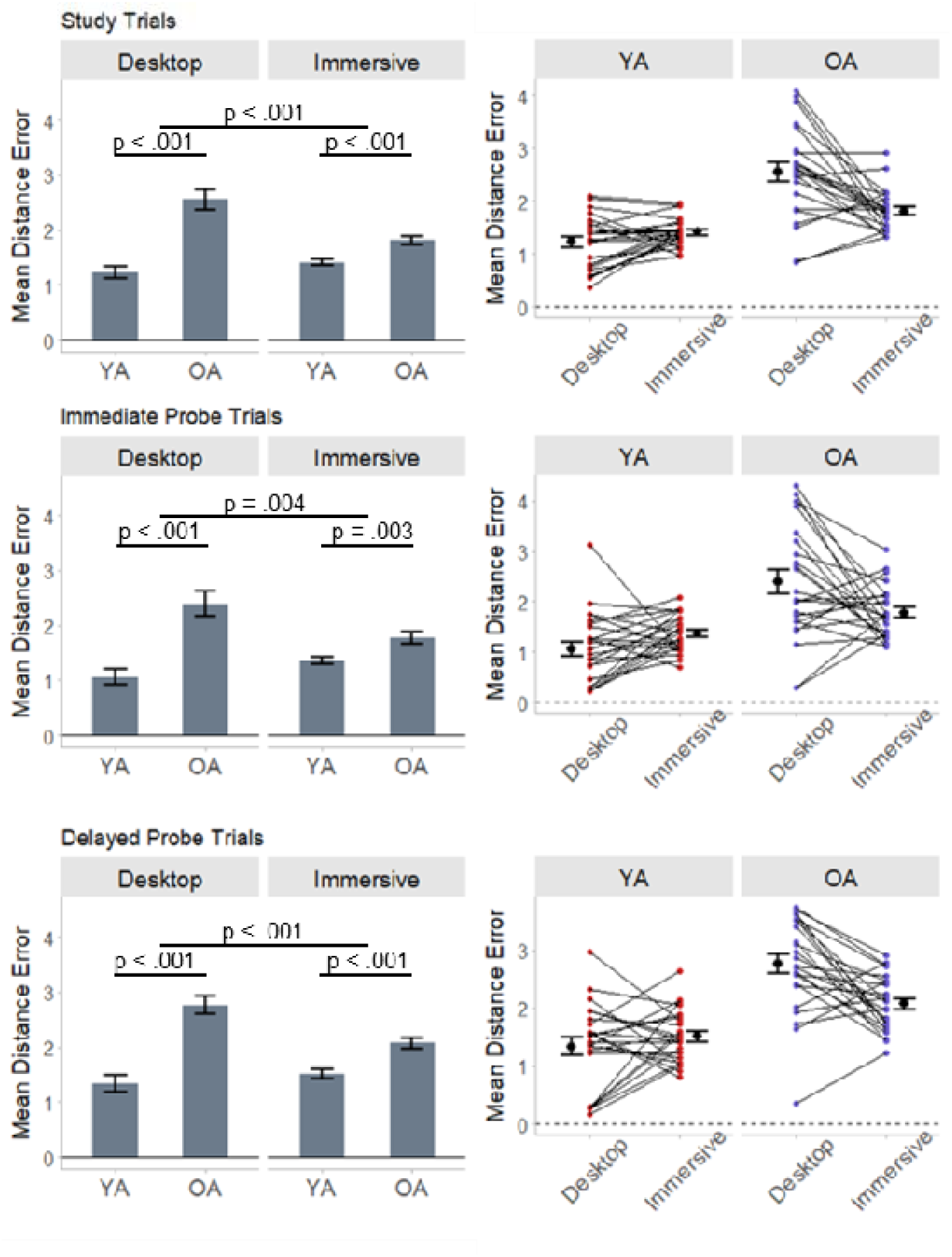
Mean spatial distance error is plotted for the spatial learning trials (top panels), immediate probe trials (middle panels), and delayed probe trials (bottom panels). The spaghetti plots illustrate within-subject changes in distance error between the respective VR conditions. Error bars reflect standard error of the mean.

The total amount of time spent navigating during the immediate and delayed probe trials (Figure 5A) did not significantly differ between age groups (immediate: *F*_(1,46)_ = 1.15, *p* = .289, partial-_η_^2^ = .024, BF_10_ = 0.483; delayed: *F*_(1,46)_ = 1.21, *p* = .277, partial-_η_^2^ = .026, BF_10_ = 0.472) or VR conditions (immediate: *F*_(1,46)_ = 0.96, *p* = .332, partial-_η_^2^ = .020, BF_10_ = 0.310; delayed: *F*_(1,46)_ = 1.94, *p* = .170, partial-_η_^2^ = .041, BF_10_ = 0.483). A significant interaction between age group and VR condition on total time spent navigating during the immediate probe trials (*F*_(1,46)_ = 5.43, *p* = .024, partial-_η_^2^ = .106, BF_10_ = 2.641) was driven by younger adults taking significantly less time to navigate in the desktop compared to immersive VR condition (*t*_(46)_ = -2.34, *p* = .024, BF_10_ = 2.83). Navigation time among older adults did not significantly differ between the respective conditions (*t*_(46)_ = 0.95, *p* = .345, BF_10_ = 0.308). The interaction between age group and VR condition on total time spent navigating during the delayed probe trials was not significant (*F*_(1,46)_ = 3.04, *p* = .088, partial-_η_^2^ = .062, BF_10_ = 0.951). Critically, the interactions between age group and VR condition on distance error remained significant when controlling for total time spent navigating during the immediate (*F*_(1,46)_ = 8.57, *p* = .005, partial-_η_^2^ = .157, BF_10_ = 16.400) and delayed (*F*_(1,46)_ = 12.79, *p* < .001, partial-_η_^2^ = .218, BF_10_ = 60.610) probe trials.

**Figure 5.**
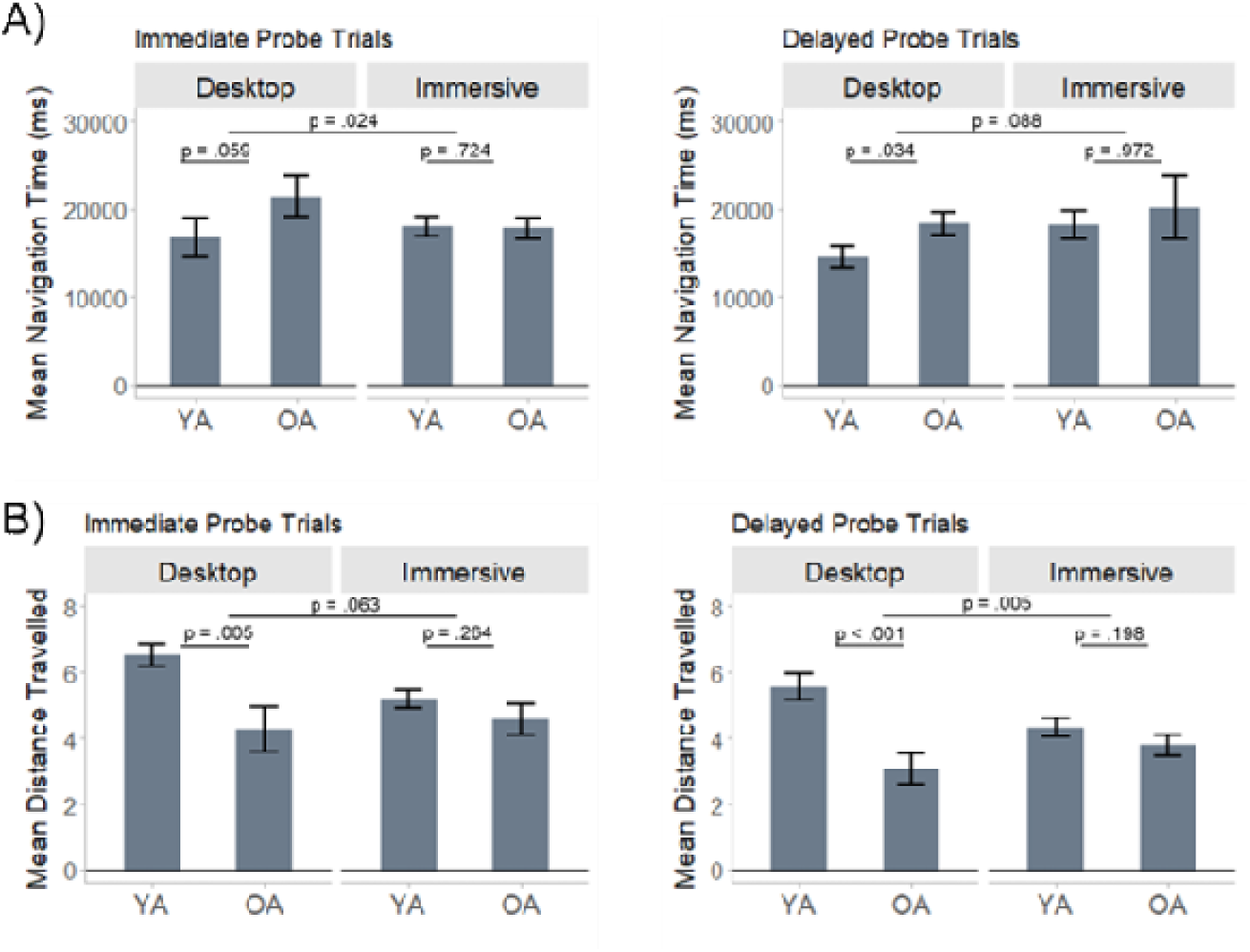
Mean (a) navigation time and (b) total distance travelled is plotted for each age group and VR condition during the immediate probe trials (left panels) and delayed probe trials (right panels). Error bars reflect standard error of the mean.

A significant interaction between age group and VR condition on the total distance travelled was evident during the delayed probe trials (*F*_(1,46)_ = 8.68, *p* = .005, partial-_η_ = .159, BF_10_ = 10.847) (Figure 5B, right panel). This interaction was driven by young adults travelling less distance in the immersive VR condition compared to the desktop condition (*t*_(46)_ = 2.66, *p* = .012, BF_10_ = 9.091). The total distance travelled by older adults on delayed probe trials did not significantly differ between the respective conditions (*t*_(46)_ = -1.50, *p* = .140, BF_10_ = 0.469). The interaction between age group and VR condition on total distance travelled during the immediate probe trials was not significant (*F*_(1,46)_ = 3.63, *p* = .063, partial-_η_ = .073, BF_10_ = 1.543) (Figure 5B, left panel). The interactions between age group and VR condition on distance error remained significant when controlling for total distance travelled on the immediate (*F*_(1,46)_ = 5.20, *p* = .027, partial-_η_^2^ = .102, BF_10_ = 3.710) and delayed (*F*_(1,46)_ = 4.62, *p* = .037, partial-_η_^2^ = .091, BF_10_ = 2.090) probe trials. As in Experiment 1, the Bayes factor revealed anecdotal evidence in favor of the alternative hypothesis after controlling for total distance travelled during the delayed probe trials.

A 2 (age group) x 2 (VR condition) ANOVA of distance error during the visible target trials revealed a significant main effect of condition (*F*_(1,46)_ = 41.89, *p* < .001, partial-_η_^2^ = .477; BF_10_ = 2887707.000) which was driven by greater distance error in the immersive relative to the desktop VR condition. The main effect of age and the age x condition interaction were not significant (*p*s > .7; BF_10_ < 0.288). A significant interaction between age group and VR condition on the time it took to reach the visible targets (*F*_(1,46)_ = 5.91, *p* = .019, partial-_η_^2^ = .114; BF_10_ = 3.563) was driven by older adults taking significantly longer in the desktop compared to the immersive VR condition (*t*_(46)_ = 3.01, *p* = .004, BF_10_ = 27.584). Time to reach the visible target did not significantly differ between the respective VR conditions in young adults (*t*_(46)_ = -0.43, *p* = .672, BF_10_ = 0.229).

Finally, we performed a 2 (experiment) x 2 (age group) x 2 (VR condition) ANOVA to examine whether spatial precision was affected by the inclusion of the additional boundary cue in Experiment 2. We observed significant main effects of experiment on spatial precision during the learning trials (*F*_(1,84)_ = 49.91, *p* < .001, partial-_η_^2^ = .363, BF_10_ = 4112.555), immediate probe trials (*F*_(1,84)_ = 27.76, *p* < .001, partial-_η_^2^ = .248, BF_10_ = 417.647), and delayed probe trials (*F*_(1,84)_ = 35.42, *p* < .001, partial-_η_^2^ = .297, BF_10_ = 2062.436). As can be seen in Figure 6, main effects of experiment were driven by reduced distance error in Experiment 2 compared to Experiment 1 (learning: *t*_(84)_ = 6.92, *p* < .001, BF_10_ = 380722.300; immediate: *t*_(84)_ = 5.27, *p* < .001, BF_10_ = 6463.548; delayed: *t*_(84)_ = 5.95, *p* < .001, BF_10_ = 194886.500). All two and three-way interactions involving the experiment were not significant (all *p*s > .14). In a follow-up ANCOVA, we included age as a covariate to rule out the possibility that reduced distance error in Experiment 2 was driven by age differences between the respective experiments (see Table 1). Main effects of experiment on distance error remained significant across all task phases (*p*s < .001). All two and three-way interactions involving the experiment also remained not significant (all *p*s > .11).

**Table 1.**
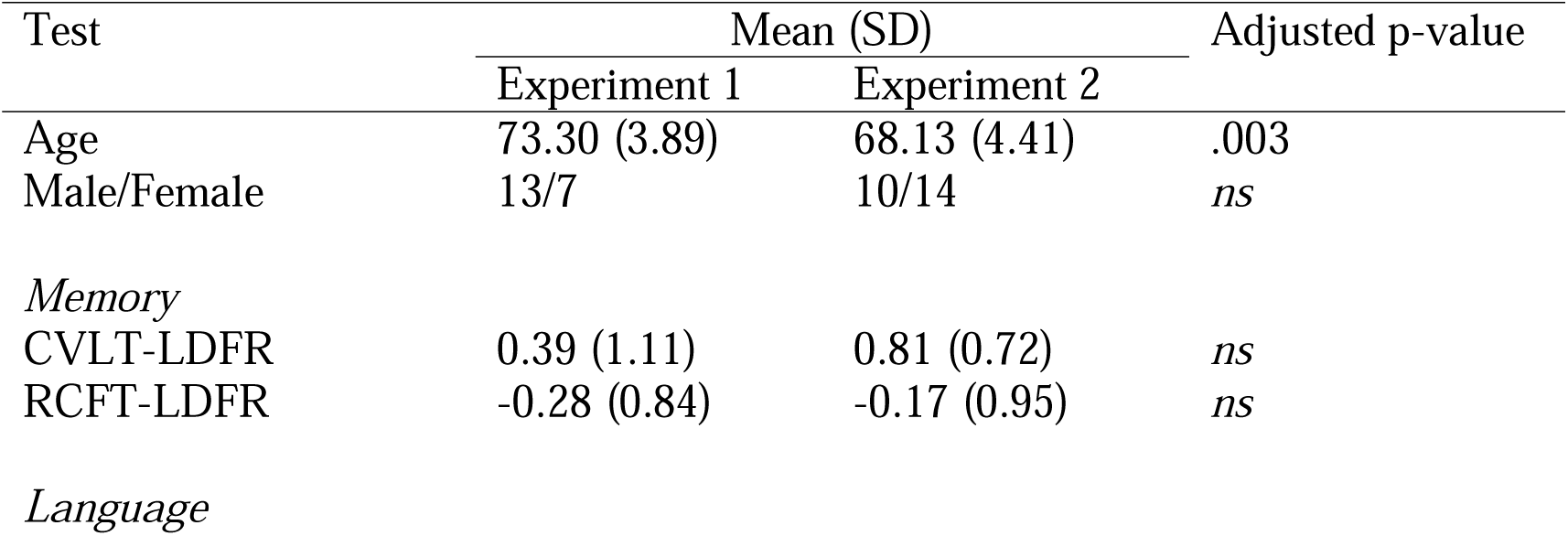
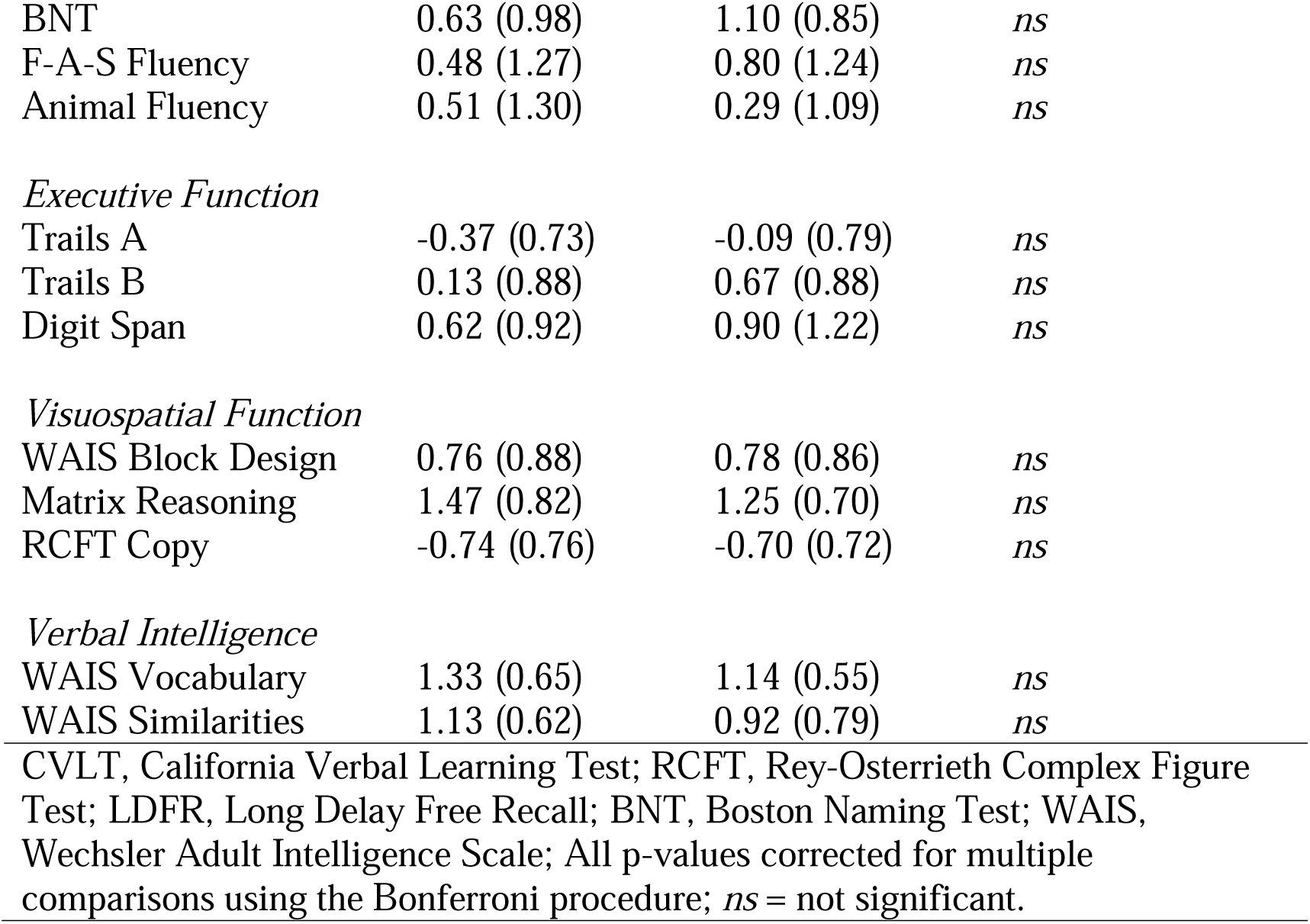
Demographics and mean (with SD) z-score performance on the neuropsychological tests for older adults.

| Test | Mean (SD) |  | Adjusted p-value |
| --- | --- | --- | --- |
|  | Experiment 1 | Experiment 2 |  |
| Age | 73.30 (3.89) | 68.13 (4.41) | .003 |
| Male/Female | 13/7 | 10/14 | *ns* |
| *Memory* |  |  |  |
| CVLT-LDFR | 0.39 (1.11) | 0.81 (0.72) | *ns* |
| RCFT-LDFR | -0.28 (0.84) | -0.17 (0.95) | *ns* |
| *Language* |  |  |  |
| BNT | 0.63 (0.98) | 1.10 (0.85) | *ns* |
| F-A-S Fluency | 0.48 (1.27) | 0.80 (1.24) | *ns* |
| Animal Fluency | 0.51 (1.30) | 0.29 (1.09) | *ns* |
| *Executive Function* |  |  |  |
| Trails A | -0.37 (0.73) | -0.09 (0.79) | *ns* |
| Trails B | 0.13 (0.88) | 0.67 (0.88) | *ns* |
| Digit Span | 0.62 (0.92) | 0.90 (1.22) | *ns* |
| *Visuospatial Function* |  |  |  |
| WAIS Block Design | 0.76 (0.88) | 0.78 (0.86) | *ns* |
| Matrix Reasoning | 1.47 (0.82) | 1.25 (0.70) | *ns* |
| RCFT Copy | -0.74 (0.76) | -0.70 (0.72) | *ns* |
| *Verbal Intelligence* |  |  |  |
| WAIS Vocabulary | 1.33 (0.65) | 1.14 (0.55) | *ns* |
| WAIS Similarities | 1.13 (0.62) | 0.92 (0.79) | *ns* |

CVLT, California Verbal Learning Test; RCFT, Rey-Osterrieth Complex Figure Test; LDFT, Long Delay Free Recall; BNT, Boston Naming Test; WAIS, Wechsler Adult Intelligence Scale; All p-values corrected for multiple comparisons using the Bonferroni procedure; *ns* = not significant.

**Table 2.**
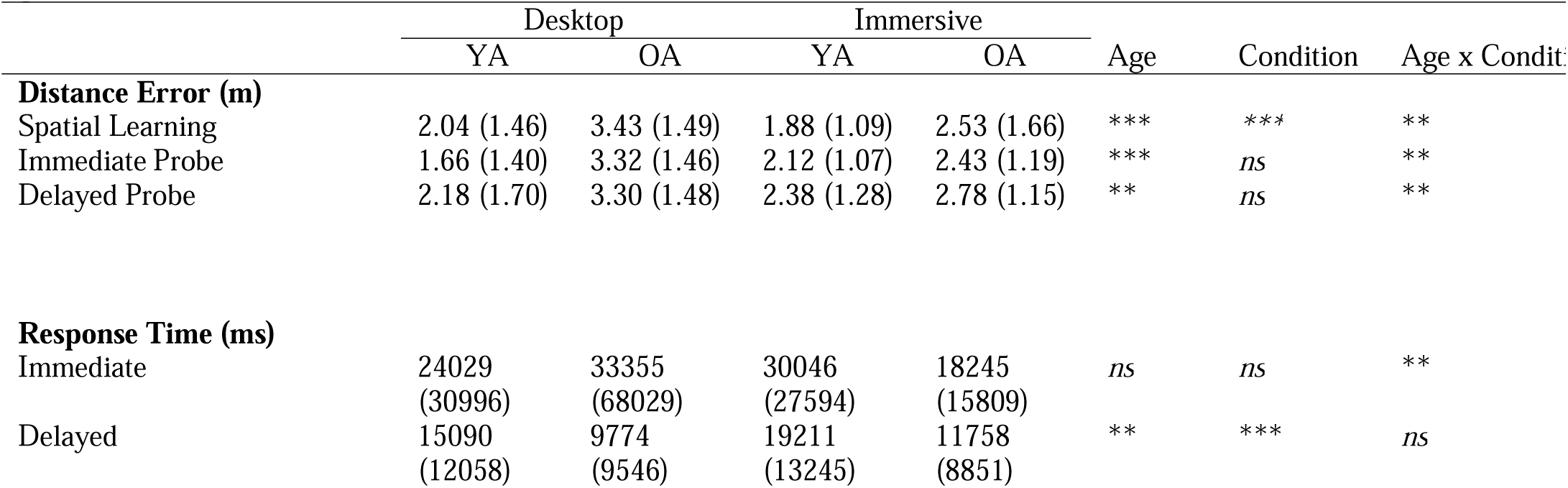
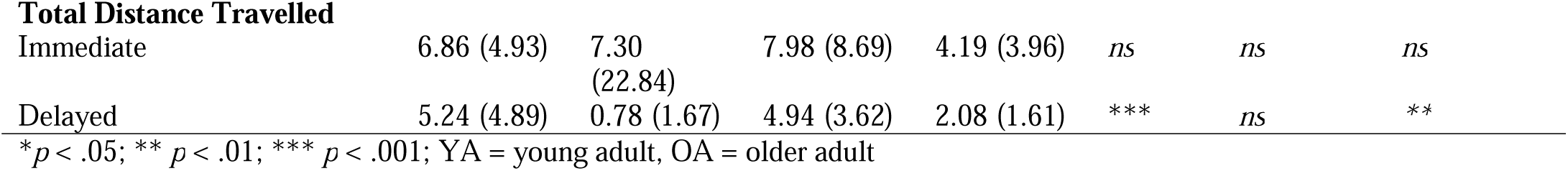
Mean (with standard deviation) of the primary outcome measures for each age group and -VR condition in Experiment 1.

|  | Desktop |  | Immersive |  | Age | Condition | Age x Condition |
| --- | --- | --- | --- | --- | --- | --- | --- |
|  | YA | OA | YA | OA |  |  |  |
| **Distance Error (m)** |  |  |  |  |  |  |  |
| Spatial Learning | 2.04 (1.46) | 3.43 (1.49) | 1.88 (1.09) | 2.53 (1.66) | *** | *** | ** |
| Immediate Probe | 1.66 (1.40) | 3.32 (1.46) | 2.12 (1.07) | 2.43 (1.19) | *** | *ns* | ** |
| Delayed Probe | 2.18 (1.70) | 3.30 (1.48) | 2.38 (1.28) | 2.78 (1.15) | ** | *ns* | ** |
| **Response Time (ms)** |  |  |  |  |  |  |  |
| Immediate | 24029<br>(30996) | 33355<br>(68029) | 30046<br>(27594) | 18245<br>(15809) | *ns* | *ns* | ** |
| Delayed | 15090<br>(12058) | 9774<br>(9546) | 19211<br>(13245) | 11758<br>(8851) | ** | *** | *ns* |
| **Total Distance Travelled** |  |  |  |  |  |  |  |
| Immediate | 6.86 (4.93) | 7.30<br>(22.84) | 7.98 (8.69) | 4.19 (3.96) | *ns* | *ns* | *ns* |
| Delayed | 5.24 (4.89) | 0.78 (1.67) | 4.94 (3.62) | 2.08 (1.61) | *** | *ns* | ** |

\* $p < .05$ ; \*\* $p < .01$ ; \*\*\* $p < .001$ ; YA = young adult, OA = older adult

**Table 3.** Mean (with standard deviation) of the primary outcome measures for each age group and VR condition in Experiment 2.

|  | Desktop |  | Immersive |  | Age | Condition | Age x Condi |
| --- | --- | --- | --- | --- | --- | --- | --- |
|  | YA | OA | YA | OA |  |  |  |
| **Distance Error (m)** |  |  |  |  |  |  |  |
| Spatial Learning | 1.25 (1.11) | 2.54 (1.51) | 1.41 (.82) | 1.81 (.93) | *** | *** | ** |
| Immediate Probe | 1.06 (.93) | 2.38 (1.45) | 1.36 (.66) | 1.78 (.85) | *** | *ns* | ** |
| Delayed Probe | 1.34 (1.24) | 2.77 (1.49) | 1.51 (.83) | 2.07 (1.00) | ** | *ns* | ** |
| **Response Time (s)** |  |  |  |  |  |  |  |
| Immediate | 16845<br>(17577) | 21667<br>(18570) | 18078<br>(6405) | 17842<br>(7097) | *ns* | *ns* | ** |
| Delayed | 14626<br>(9523) | 18420<br>(9732) | 18211<br>(10630) | 19931<br>(33084) | ** | *** | *ns* |
| **Total Distance Travelled** |  |  |  |  |  |  |  |
| Immediate | 6.53 (2.55) | 4.27 (4.62) | 5.18 (1.74) | 4.57 (3.10) | ** | *ns* | *ns* |
| Delayed | 5.55 (3.36) | 3.09 (3.00) | 4.33 (1.94) | 3.76 (2.30) | *** | *ns* | ** |

\* $p < .05$ ; \*\* $p < .01$ ; \*\*\* $p < .001$ ; YA = young adult, OA = older adult

**Figure 6.**
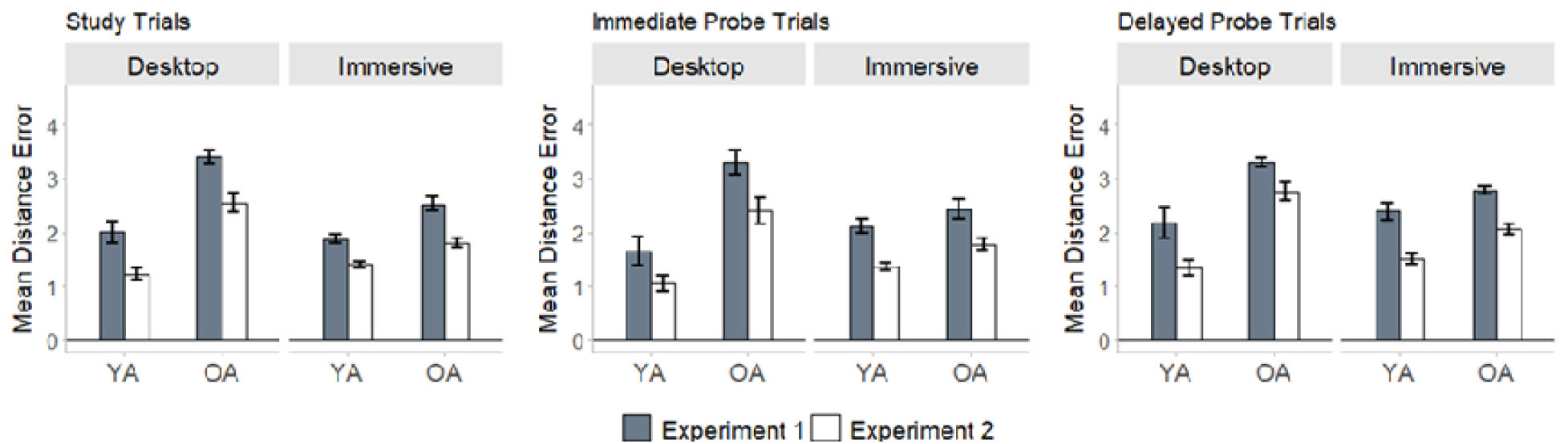
Distance error was significantly reduced in Experiment 2 compared to Experiment 1 during spatial learning, immediate probe trials, and delayed probe trials. The main effect of experiment was not moderated age group or VR condition.

### Discussion

Spatial distance error in young and older adults was reduced in Experiment 2 compared to Experiment 1. These results underscore the behavioral advantages afforded by inclusion of additional spatial boundaries to navigate, replicating numerous prior studies (Bécu et al., 2019, 2023; Ying et al., 2023). Despite this apparent advantage, however, spatial precision in older adults was reliably reduced when attempting to learn and recall spatial locations in a two-dimensional virtual desktop environment compared to immersive VR during Experiment 2. We thus replicated the principal findings from Experiment 1 in an independent sample of young and older adults. Despite an overall increase in spatial precision, the inclusion of an additional geometric boundary cue did not influence the overall effect of VR modality on age differences in spatial performance. These results therefore confirm that the principal findings from Experiment 1 could not be wholly accounted for by age-related differences in distal landmark processing.

## General Discussion

Models of human aging and navigation are largely predicated on findings from experimental paradigms employing desktop-based VR to assay spatial abilities (Lester, Moffat, Wiener, Barnes, & Wolbers, 2017; Moffat, 2009). Across two experiments, we observed robust evidence that age-related differences in the precision of spatial memories in a virtual MWM task were moderated by the modality of the virtual testing environment. Reliable age differences in immediate and delayed spatial recall were evident in a virtual desktop condition that required using a keyboard and mouse to navigate, replicating numerous prior studies (Daugherty & Raz, 2017; Daugherty et al., 2015; Iaria et al., 2009; McAvan et al., 2021; Moffat et al., 2007; Moffat & Resnick, 2002; Zhong et al., 2017). The magnitude of these age differences was attenuated, however, when tested in an immersive virtual condition that permitted free ambulation during navigation. These effects were driven by disproportionately worse performance in older adults when navigating a virtual desktop environment, which restricted self-generated idiothetic feedback, involved a more limited field of view, and was potentially less familiar and natural to the older cohort. These findings are also consistent with prior observations that age differences in path integration are reduced or even eliminated when multisensory cues are available during navigation (Adamo et al., 2012; Allen et al., 2004). We emphasize, however, that the vMWM task evokes spatial computations that are inherently distinct from those involved with successful path integration, most critically, the need to integrate visual landmarks with idiothetic-driven path integration computations (Ekstrom & Isham, 2017; Warren et al., 2001; Zhou & Mou, 2018). Our findings thus provide new insights into the nature of age differences in spatial memory by suggesting that at least one factor driving the extent of these differences relates to the testing modality.

Although age-related differences in spatial precision were significantly attenuated when tested in the immersive and ambulatory VR environment, we note that reliable age differences were evident in this condition. This is consistent with the wealth of rodent studies showing age differences when navigating in ‘real space’ (Lindner, 1997; Shen & Barnes, 1996) and a more limited literature involving older humans navigating real-world spaces (Kirasic, 1991; Kirasic, Allen, & Haggerty, 1992; Gazova et al., 2013; Newman & Kaszniak, 2000). This finding also replicates findings from a recent study from our laboratory using a similar version of the vMWM task in immersive VR (McAvan et al., 2021). Together, these findings suggest that, even under more naturalistic settings, some age-related differences in spatial precision persist. Nevertheless, we believe the findings reported in this paper underscore important limitations in using desktop VR to assay spatial navigation deficits cross sectionally.

Spatial navigation is an inherently complex construct that draws on multiple sensory systems (e.g., visual, proprioceptive, vestibular, and motoric), each of which is susceptible to advancing age (Agrawal et al., 2012; Gadkaree et al., 2016; Shaffer & Harrison, 2007). Our findings suggest that age differences in the precision of spatial memories are significantly reduced in an immersive virtual environment in which visual and self-motion cues are available to guide navigation. One possibility is that older adults place a greater reliance on complementary sensory cues to compensate for reduced acuity in other sensory domains. Consequently, the lack of body-based cues in the desktop environment, which favored primarily visual input, may have affected older adults’ ability to encode spatial representations in sufficient detail to support accurate spatial memory (Ekstrom & Hill, 2023). Though this account is speculative, it is consistent with prior reports that age differences in navigation performance arise, at least in part, from a failure to combine information from different sensory modalities, resulting in noisier spatial representations (Bates & Wolbers, 2014; Mahmood et al., 2009; Merhav & Wolbers, 2019).

Notably, the desktop and immersive VR environments differed across several dimensions which may affect observed age differences in spatial performance. For example, older adults may have uniquely benefited from the expanded three-dimensional field of view in the immersive VR condition (Tan et al., 2006). Likewise, navigation performance in desktop VR may be confounded by age differences in computer fluency and prior gaming experience (Charness & Boot, 2022). Most older adults have less experience with video games, in part because gaming is still largely geared and marketed for a much younger audience (Gamberini et al., 2006; Salmon et al., 2017). Experience with computer games, which involves how to combine use of a joystick with the movement of optic flow on a desktop computer screen, however, is crucial to learning how to navigate in virtual reality. Reconciling the limitations of desktop-based VR, especially in cross-sectional research, is critical for understanding the biological significance of prior findings derived from MRI and mobile-based gaming apps (e.g., Coutrot et al., 2018).

In the foregoing discussion, we suggest that impaired spatial memory observed in older adults navigating a desktop environment stem from a lack of self-generated idiothetic feedback and/or a combination of age-related confounds such as limited exposure to computer gaming equipment or reduced visual and sensorimotor acuity. Although these possibilities are certainly not mutually exclusive, the present data do not allow us to adjudicate between the factors that drove spatial memory deficits in the desktop VR environment. Another limitation of the present study concerns the ecological validity of the sparse environment encountered in the vMWM task. Prior work has shown that age differences in navigation performance are attenuated when navigating in highly familiar environments, such as a local supermarket (Kirasic, 1991). Future research is needed to address whether the moderating effect of VR modality extends to familiar and/or hyper-realistic environments that more closely approximate the types of scenes and challenges older adults face in their daily lives.

In conclusion, marked age differences in the precision of spatial memories were evident when using a keyboard and mouse to navigate a virtual desktop MWM environment. These age differences were reduced, however, when navigating in a fully immersive and ambulatory vMWM environment that permitted unrestricted locomotion and reduced age-related confounds associated with navigating a virtual desktop environment. These results underscore the importance of developing naturalistic and ecologically valid assays of spatial behavior, especially for cross-sectional studies comparing spatial performance in young and older individuals.

## Acknowledgements

De-identified data and analysis scripts are posted on https://osf.io/5vbzn/. A non-peer reviewed preprint of this manuscript is available on bioRxiv https://doi.org/10.1101/2023.01.23.525279. The authors wish to thank Stephanie Doner, Kaela Liddle, Kate Carpenter, Erin Rose, Makena Lees, Destiny Sanchez, Delaney Sanders, and Don Markham for their assistance with data collection. The authors would also like to thank the research volunteers who dedicated their time to make this research possible.

